# Estimating the distance to an epidemic threshold

**DOI:** 10.1101/247734

**Authors:** Eamon B. O’Dea, Andrew W. Park, John M. Drake

## Abstract

The epidemic threshold of the susceptible-infected-recovered (SIR) model is a boundary separating parameters that can permit epidemics from those that cannot. This threshold corresponds to points where the stability of the system’s equilibrium reaches zero. Consequently, we use the average rate at which deviations from the equilibrium shrink to define a distance to this threshold. However, the vital dynamics of the host population may occur slowly even when transmission is far from threshold levels. Here we show analytically how such slow dynamics can prevent estimation of the distance to the threshold for some individual variables of the model. Although these results are exact only in the limit of long-term observation of a large system, we find that they still provide useful insight into the behaviour of estimates from simulations with a range of population sizes, environmental noise, and observation schemes. Having established some guidelines about when estimates are accurate, we then illustrate how multiple distance estimates can be used to estimate the rate of approach to the threshold. The estimation approach is general and may be applicable to zoonotic pathogens such as MERS-CoV as well as vaccine-preventable diseases such as measles.

## Introduction

Many infectious disease epidemics occur with sufficient regularity that their anticipation is straightforward. For example, seasonal influenza has a pronounced winter seasonality in most of the world, with annual outbreaks [1]. Some systems are more episodic but still well-understood, such as measles in sub-Saharan Africa where regional inter-epidemic periods between 1–4 years have been observed in recent times [2]. In contrast, emerging and re-emerging infectious diseases are rarely anticipated, even though the root causes are often discerned soon after the event. Many childhood infectious diseases naturally spread effectively, including measles, chickenpox and rubella. This means that in unvaccinated populations, one infectious individual may infect many others, measured by the pathogen’s basic reproduction number, *R*_0_ [3]. Outbreaks are prevented in these cases by maintaining a very high proportion of vaccinated individuals, generating herd immunity in which the effective reproduction number is below 1, meaning small chains of transmission are quickly broken [4]. Reduced vaccine uptake rates can move the infectious disease system from controlled (sub-critical, with effective *R*_0_ < 1) to super-critical when outbreaks may occur [5]. Alternatively, other features of the system may be slowly changing, similarly enhancing the transmission of the pathogen. Host demographic changes, particularly rising birth rates, can increase the supply of susceptible individuals to the population, and pathogens frequently evolve at high rates, whereby fitter strains (higher *R*_0_) may be favoured by selection [6]. Predicting a dynamical system’s movement from sub- to super-critical before it happens has enormous potential to remove the element of surprise associated with emerging infectious diseases, to prioritise mitigation strategies to reverse, stop, or slow the transition, and in worst cases to simply be better prepared for the inevitable. Recent work has also illustrated that following a transition from sub- to super-critical there is a characterisable bifurcation delay—a waiting time until the outbreak actually occurs following suitable conditions being met [7]. Consequently, estimates of how far a system is from the epidemic threshold could help public health officials make judgements about policy, infer on which side of the threshold the population lies, and track the movement of a system towards a threshold (providing early warnings) and even away from a threshold as a way of evaluating the effectiveness of any external changes to the system aimed at controlling infectious disease outbreaks.

A potentially robust basis for estimating the distance to a threshold is the general slowing down of a system’s dynamics as a threshold is approached. To be more precise, the average decay rate of deviations from a fixed point of the system becomes increasingly smaller as the parameters of the system approach the point at which that fixed point becomes unstable. Wissel [8] pointed out that this phenomenon, known as *critical slowing down* or sometimes simply as *slowing down*, could be used to determine whether the parameters of a system were approaching a threshold that, when crossed, could result in the system changing in an abrupt and drastic manner. Such changes have come to be called *critical transitions* [9]. Recently a great deal of interest has developed in the possibility of devising model-independent methods to anticipate critical transitions in complex systems using early-warning signals [10]. In general, early-warning signals are statistical properties of observations of systems that can be expected to change in characteristic ways as a threshold is approached. Perhaps the most common examples are increasing autocorrelation and variance of model variables. These signals can often be derived from the increasingly slow decay of perturbations due to slowing down, and many other early-warning signals are in one way or another quantifications of slowing down. The beauty of early-warning signals is that their basis in generic properties of dynamical systems means they have the potential to be reliable even when the system is complex and unidentifiable. Examples of complex and poorly identified systems abound in ecology and epidemiology. With application to such systems in mind, several authors [11, 12, 13] demonstrated the application of early-warning signals based on slowing down to forecasting infectious disease emergence and eradication. Further development and integration of these methods into surveillance systems may provide a novel and broadly applicable method of evaluating the control of infectious diseases from existing surveillance data streams.

To explain some of the current challenges in further developing approaches to estimating the distance to the threshold, we will make reference to some elements of dynamical systems theory. Following Wiggins [14], a general dynamical system may be written as a system of equations for a vector field *ẋ* = *f*(*x, θ*), where the overdot indicates a derivative with respect to time, *x* is a vector of real numbers that determine the point of the system in its phase space, and *θ* is a vector of real numbers that are parameters of the system. A solution to the system is a function *x* of time that over some time interval satisfies *ẋ* = *f*(*x*(*t*), *θ*). A fixed point *x*\* of the system is a solution that does not change with time (*i.e*., it satisfies 0 = *f*(*x*\*, *θ*)). Such a point is also referred to as a steady state or an equilibrium of the system. A fixed point is called asymptotically stable if solutions that start at points near the fixed point move closer to it over time. Because the starting points are nearby, deviations *z* = *x* – *x*\* are small and can be accurately modelled by solutions to the linear system *ż* = *Fz*, where *F* denotes the matrix of first derivatives of *f* with respect to *x* (*i.e*., the Jacobian matrix). The general solution of such a system is *z*(*t*) = exp(*Ft*)*z*(0). If the real parts of all of the eigenvalues of *F* are negative, this solution will shrink to zero and it follows that *x*\* is asympotically stable. If the real parts of any of the eigenvalues are positive, the solution will not shrink to zero and *x*\* is not asymptotically stable. Thus, as long as the real parts of the eigenvalues of *F* are not zero, their signs tell us whether or not any fixed point is stable.

The relationship between the speed of a system’s dynamics and the distance to the threshold arises in the common case that the eigenvalues of *F* are continuous functions of the parameters *θ* of the system and none of the eigenvalues have zero real parts. In this case for a stable fixed point to become unstable, one of the eigenvalues must cross zero. Thus as the parameters approach the threshold where stability is lost, one of the eigenvalues must approach zero in its real part. We call such an eigenvalue an *informative eigenvalue* since its value is informative of how far the system’s parameters are from a threshold. We call the magnitude of such an eigenvalue a distance to the threshold. If an informative eigenvalue can be monitored over time, one can determine whether the system is approaching a threshold or not and even make a forecast of when the threshold will be crossed. An informative eigenvalue can be measured by monitoring the decay of small perturbations away from the fixed point along the eigendirection of the informative eigenvalue. Identifying trends in such a decay rate is the goal of early-warning signals based on slowing down.

Despite the simplicity of this goal, it is currently not clear exactly how it can be achieved when systems have multi-dimensional phase space. When one of the eigenvalues of *F* gets closer to zero, only a small number of the model’s observable variables may become less resilient to perturbations. The implication is that early-warning signals such as increasing variance and autocorrelation will not be present in all of the model’s variables. Several authors have provided examples of such a case. Kuehn [15] showed that in a susceptible–infected–susceptible (SIS) model of an epidemic on an adaptive contact network, only one of the three model variables had a clear increase in variance as the epidemic threshold was crossed. Boerlijst *et al*. [16] even showed that, depending on the types of perturbations a system experiences, the autocorrelation of some variables may either increase or decrease as a threshold is approached. Consequently, a recent review [17] identified the selection of appropriate variables in multivariate systems for detection of slowing down as an important problem in need of solution. Dakos [18] has recently used an eigendecomposition of *F* to derive a simple rule about which state variables have a decay rate that is most affected by the dominant eigenvalue of *F*. However, this approach only provides a partial answer to the question of variable selection because it does not account for the covariance of the perturbations to the system, which can be as important as the eigenvectors of *F* on the decay rate of a state variable. Furthermore, another consequence of models having multiple dimensions is that the informative eigenvalue may not necessarily be the dominant eigenvalue. When its real part gets close enough to zero, the informative eigenvalue will of course become dominant but, as we shall demonstrate, that may not happen until it is very small. So although slowing down is often explained to be a consequence of the dominant eigenvalue approaching zero, methods to estimate the dominant eigenvalue of *F* from a multivariate time series may not reliably estimate the distance to the threshold. There does not seem to be any general approach for estimating the distance to the threshold in multidimensional systems.

In this work, we derive an explicit relationship between the eigenvalues of *F* and the autocorrelation function of each of the variables in a multivariate system. The resulting equations lead us to a simple condition for determining the types of perturbations under which estimation of a variable’s autocovariance function can be translated into an estimate of the distance to the threshold. We demonstrate the application of this method to the susceptible–infected–removed (SIR) model for directly transmitted infectious diseases. We find that for parameters relevant to many vaccine-preventable diseases, the autocorrelation of the number infected almost always is indicative of the distance to the epidemic threshold, while the autocorrelation of the number susceptible is not. We examine the sensitivity of the accuracy of these estimates to environmental noise, small population size, the frequency of observation, and observation of case reports instead of the actual number infected. We also show a simple example of estimating the change in the distance to the threshold over the length of a time series. These results demonstrate the general feasibility of developing statistical systems for forecasting disease emergence and documenting the approach to elimination.

## Methods

### Model

The model that motivated the development of the following methods is the susceptible–infected–removed (SIR) model with demography. We let *X*(*t*) denote the number of susceptible individuals, *Y* (*t*) the number of infected (and infectious) individuals, *Z*(*t*) the number of removed individuals (recovered or vaccinated), and *N*(*t*) = *X*(*t*) + *Y* (*t*) + *Z*(*t*) the total population size at time *t*. Typically, we assume that these numbers are the integer-valued random variables of a Markov process having the parameters defined in table 1 and the transitions defined in table 2. In the following, we often omit explicit notation of time dependence for the sake of conciseness. We also consider models where the death rate or the force of infection (*i.e*., the per capita rate at which susceptibles become infected) is subject to variation over time due to fluctuations in the environment over time. We follow Bretó and Ionides [19] in modelling such variation as multiplicative gamma white (“temporally uncorrelated”) noise. This noise could represent changes in rates due to weather conditions or social mixing [20] or even model errors in model specification, such as a failure to model spatial heterogeneity [21]. Bretó and Ionides [19] show that the model remains Markovian with such noise with the modified propensities for the death and transmission events given by the expressions in table 3. Inclusion of multiplicative gamma noise leads to the possibility that more than one individual becomes infected or dies in a single event (*i.e*., the associated counting processes are compound) and table 3 gives the propensity of births and death events for all positive integers *k, k*_1_, *k*_2_ and *k*_3_.

**Table 1:** Model parameters

| Symbol | Definition | Default value |
| --- | --- | --- |
| $\eta$ | importation rate | $1/(3\sqrt{N_0}) \text{ year}^{-1}$ |
| $\beta$ | transmission rate | varied |
| $\gamma$ | recovery rate | $365/22 \text{ year}^{-1}$ |
| $\mu$ | death rate | $0.02 \text{ year}^{-1}$ |
| $N_0$ | initial population size | $10^7$ individuals |
| | observation frequency | $52 \text{ year}^{-1}$ |
| $\tau_f$ | magnitude of environmental noise in force of infection | 0 |
| $\tau_d$ | magnitude of environmental noise in death rate | 0 |

**Table 2:** Transitions of the SIR model

| Name | $(\Delta X, \Delta Y, \Delta Z)$ | Propensity |
| --- | --- | --- |
| birth | ( 1, 0, 0) | $N_0\mu$ |
| death of $X$ | (-1, 0, 0) | $X\mu$ |
| death of $Y$ | ( 0, -1, 0) | $Y\mu$ |
| death of $Z$ | ( 0, 0, -1) | $Z\mu$ |
| transmission | (-1, 1, 0) | $XY\beta/N_0 + X\eta$ |
| recovery | ( 0, -1, 1) | $Y\gamma$ |

**Table 3:**
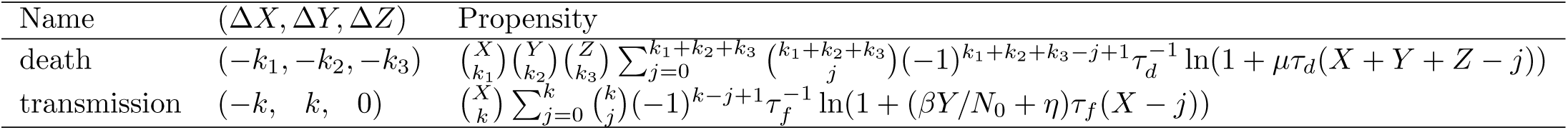
Modified transitions of the SIR model allowing for environmental heterogeneity Name

| Name | $(\Delta X, \Delta Y, \Delta Z)$ | Propensity |
| --- | --- | --- |
| death | $(-k_1, -k_2, -k_3)$ | $\binom{X}{k_1} \binom{Y}{k_2} \binom{Z}{k_3} \sum_{j=0}^{k_1+k_2+k_3} \binom{k_1+k_2+k_3}{j} (-1)^{k_1+k_2+k_3-j+1} \tau_d^{-1} \ln(1 + \mu\tau_d(X+Y+Z-j))$ |
| transmission | $(-k, k, 0)$ | $\binom{X}{k} \sum_{j=0}^k \binom{k}{j} (-1)^{k-j+1} \tau_f^{-1} \ln(1 + (\beta Y/N_0 + \eta)\tau_f(X-j))$ |

There are several biological assumptions implicit in our model. We use the standard assumption of frequency-dependent transmission, which has been shown to be a more appropriate model than the common alternative assumption of density-dependent transmission for a number of infectious diseases [22]. However, we calculate frequency using the parameter *N*_0_ instead of *N*(*t*). The initial population size *N*_0_ is also the expected value of the population size because we set the birth rate in table 2 as *N*_0_*μ*. Another assumption is that the average death rate of individuals is constant throughout their lifetimes. According to Anderson and May [23], this is a common assumption among the traditional literature in mathematical epidemiology. It is biologically accurate in that humans are subject to a small and relatively constant mortality rate until they reach old age. A more realistic model would include much higher mortality at old age, but such realism is not necessary for our study. Another key feature of our model is the inclusion of the *η* term in the force of infection (tables 2 and 3), which relaxes the assumption that the population is closed to infection from other populations or environmental reservoirs. We include such a term to allow our model to represent populations in which an infectious disease is repeatedly introduced but unable to persist within the population.

Although the model is stochastic, the expected value of the model’s variables is deterministic. The rate of change in the expected value when the system is in a given state can be approximated summing over all possible updates in tables 2 and 3 and weighting each update by its propensity [24]. Calculating the rate of change in the expected value of *X*, *Y*, and *Z* in this way leads to the following system of differential equations

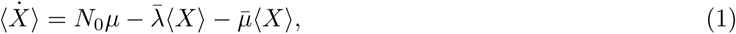

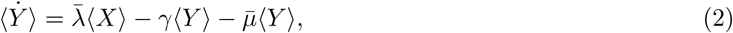

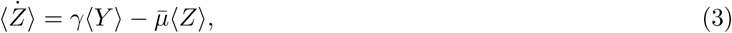

where the overdot indicate a time derivative and where

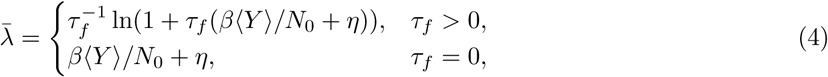

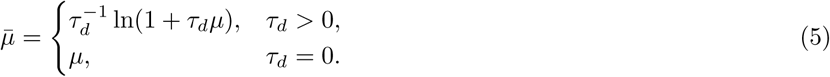

The equations for 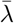 and 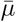 in the case of non-zero environmental noise are the infinitesimal means derived in Bretó and Ionides [19]. By setting these equations equal to zero and solving for 〈*X*〉 and 〈*Y*〉, we can find the approximate fixed point of the system for a given set of model parameters.

The equations for the fixed point of the differential equations allow us to explain what we mean by epidemic threshold. For the sake of clarity, we consider the equations only when *τ_d_* and *τ_f_* are zero. In that case the exact equation for the *Y*-coordinate of the fixed point, which we denote *Y*\*, is

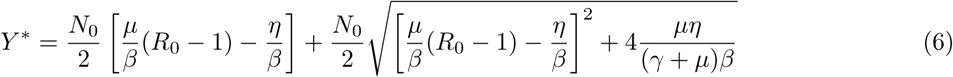

where *R*_0_ = *β*/(*γ*+*μ*). *R*_0_ is known as the basic reproduction number and we consider the epidemic threshold for the SIR model to be the surface in parameter space where *R*_0_ = 1 and *η* = 0. To see why, note that when *η* = 0, equation (6) has a non-zero value only when *R*_0_ > 1; only when *R*_0_ > 1 will the introduction of an infection into a susceptible population lead to an epidemic according to the system of differential equations. Accordingly, one can interpret *R*_0_ as the average number of new infections caused by an infected individual in a susceptible population. From the point of view of fixed points, the epidemic threshold separates the region of parameter space where a fixed point occurs with *Y*\* = 0, a disease-free equilibrium, from the region where a fixed point occurs with some *Y*\* > 0, an endemic equilibrium. In short, we define the epidemic threshold for the SIR model as the location of the model’s bifurcation. In the electronic supplementary material, we show how this definition remains useful in the case that *η* > 0 and the bifurcation is imperfect.

For other more realistic models, such as the structured immunity model of Reluga *et al*. [25], there may be multiple stable states for a given value of the parameters. For such models, there may not be a single bifurcation occurring at *R*_0_ = 1 but instead multiple bifurcations at different points in parameter space. The term epidemic threshold is ambiguous for such models because, for example, the fraction of the population infected may jump up as the parameters cross one threshold but not jump down until the parameters moved much farther backward in the opposite direction. However, our methods are applicable to estimating the distance to any threshold parameter values that corresponds to the loss of an equilibrium’s stability.

### Relating eigenvalues to autocovariance

When the dynamics are characterised by small fluctuations around a fixed point, the degree of autocorrelation of these fluctuations may be indicative of the distance to the threshold. A first step in demonstrating this relationship is to derive a probability density function for the fluctuations. Let *z*(*t*) denote a vector of deviations from the fixed point that is in units of the square root of the system’s size. Let *p*(*z*) be the probability density function of these deviations. In the limit of a large system size, this function may be approximated as the solution to the Fokker–Planck equation

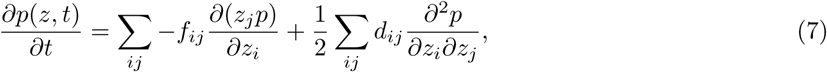

where the matrix *F* (with elements *f_ij_*) determines the expected trajectory of *z* toward zero and the matrix *D* (with elements *d_ij_*) describes the covariance of a Gaussian white noise process that acts on *z*. The matrices *F* and *D* follow directly from the transition probabilities. For the SIR model in the previous subsection, we take *N*_0_ as the system size, 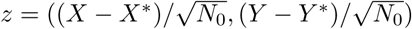 and obtain

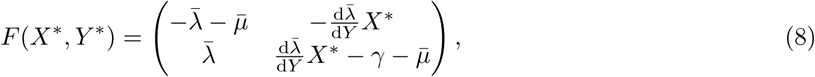

where 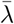 and 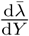 are evaluated at *Y* = *Y*\*. Note that we have omitted deviations from *Z*\* in our vector *z*. Including these deviations would be straightforward but the behaviour of *Z* is in many ways similar to that of *X* and the value of *Z* does not affect the rates at which *X* and *Y* change (table 2). Therefore, we have omitted the fluctuations of *Z* in the following to make our results more concise. For the covariance matrix, we obtain

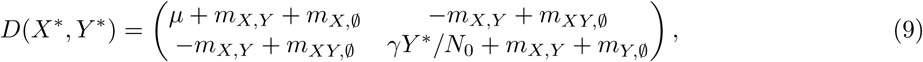

where

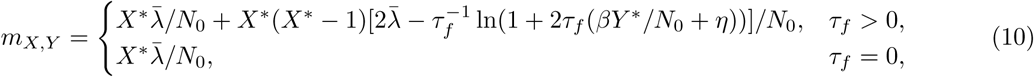

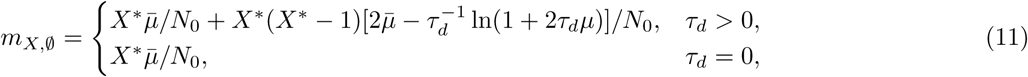

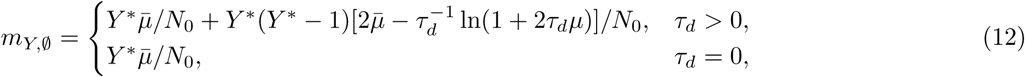

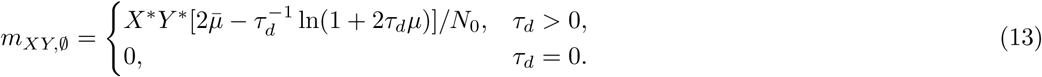

A solution to equation (7) is a Gaussian density function with a mean of zero and a covariance matrix Σ (with elements *σ_ij_*) that depends on *F* and *D*. van Kampen [24] provides a detailed introduction to these methods.

For these Gaussian solutions, the autocovariance function of the deviations may be written in terms of the eigenvalues of *F*. The relationship is particularly simple when the eigenvectors of *F* are used as the basis of the coordinates. Thus let 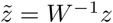, where *W* is a matrix of the eigenvectors of *F*, and let 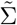, denote the covariance matrix of 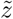. Then, using the decomposition of Kwon *et al*. [26], it follows that

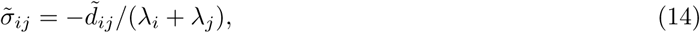

where *λ_i_* denotes an eigenvalue of *F* and we assume that all of these eigenvalues are distinct. The autocovariance matrix is defined as 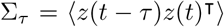, where the angular brackets denote expected value over time or realisations of the system. It follows from the stationarity of the solution that 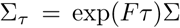. In the eigenvector basis, we have 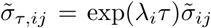. Thus the behaviour of the autocovariance along an eigendirection as a function of the lag *τ* is a simple and identifiable function of the corresponding eigenvalue. If *λ_i_* is real, then 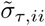 decays exponentially toward zero at the rate *λ_i_*. If *λ_i_* has an imaginary component, then the real and imaginary parts of 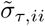 oscillate around zero with a frequency given by the imaginary component of *λ_i_* and an amplitude that decays exponentially at the rate given by the real component of *λ_i_*. Since 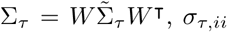 will be a linear combination of the elements of 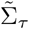 (see equation S4 in the electronic supplement). Therefore, the elements of the autocovariance matrix Σ*_τ_* are linear combinations of functions from which the eigenvalues of *F* are identifiable.

### Solving for the space of suitable noise parameters

The relationship between the eigenvalues and the autocovariance established in the previous subsection clarifies the question of when the autocovariance of a variable contains sufficient information to estimate the distance to a threshold. Any threshold corresponds to an eigenvalue crossing zero. Recall that we call such an eigenvalue an informative eigenvalue and that the magnitude of its real part can be considered the distance to the threshold. If it is known that the imaginary part of the eigenvalue will also be zero at the threshold, then the magnitude of the imaginary part can be considered a second component of the distance. Note that in the case that an informative eigenvalue is complex it will be a part of a conjugate pair. Estimation of the decay rate and frequency of oscillation of a variable’s autocovariance function can provide an estimate of the distance to the threshold when they are close to the real and imaginary parts of an informative eigenvalue. This condition on the autocovariance function for an estimate to be accurate, together with equations 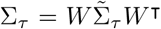 and equation (14), can be translated into conditions on the eigenvectors *W* of *F* and the covariance matrix *D* of the perturbations. Thus, we now have a general link between the parameters of models and the potential for a model variable to provide an estimate of the distance to the threshold. In the electronic supplementary material, we provide an explicit calculation of the values of *D* that permit a distance estimate for each variable.

### Obtaining distance estimates from time series

We estimate the distance to the threshold from a time series as follows. The main idea is to suppose that the autocorrelation will exponentially decay with increasing lags at a rate equal to the real part of the informative eigenvalue and that any oscillations in the autocorrelation function have a frequency equal in magnitude to the imaginary part of the informative eigenvalue. The first step is then to estimate the autocorrelation of the time series for a series of lags, which we did using the acf function in R. Because sometimes the autocorrelation can have cycles with a period of several years, we used lags from 0 to 30 observations less than the length of the time series. Next we use a nonlinear least-squares optimiser to fit two models for the estimated autocorrelation 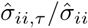:

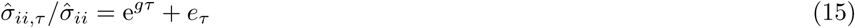

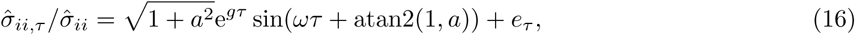

where *e_τ_* is an error term, *g* is the decay rate parameter, *ω* is the frequency parameter, and *a* is a phase angle parameter, and atan2 is the inverse tangent function with arguments in the order *y, x*. We use the nlsLM function to fit these models. This function is available in the minpack.lm package [27], and it provides an R interface to the Levenberg–Marquardt optimiser in the MINPACK library. We used nlsLM instead of the nls function that comes with R because it was less sensitive to the choice of initial values of the parameters for the optimisation of the model fit. For initial values, we set *a* to zero, *g* to the least-squares slope of the log of the absolute value of the estimated autocorrelation versus the lag, and *ω* to the frequency that maximised the spectral density of the estimated autocorrelation. Only autocorrelations with relatively small lags were used for the initial estimate of *g*. Specifically, all lags including and following the first lag that was less in magnitude than 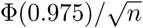 where Ф is the cumulative distribution function of a standard normal random variable and *n* is the length of the time series. We fit the data with and without an oscillation component in the model and used the following information criteria to evaluate the models:

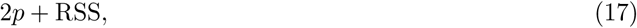

where RSS stands for residual sum of squares, *p* = 1 for model (15), and *p* = 3 for model (16). We used the estimates from the model with the lower score. If neither model’s information criteria exceeded 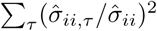, we concluded that the data contained insufficient information to provide an estimate. When estimates were available, we calculated the distance to the threshold as 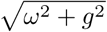.

If the input time series consisted of aggregated counts, we modified the model having no oscillations to

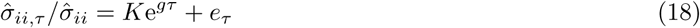

and we excluded the lag-0 autocorrelation from our data to be fitted. This model, which has a new parameter *K* that determines the autocorrelation at a lag of 1, matches the form of the autocorrelation function for counts of deaths in a birth-death-immigration model [28]. The birth-death-immigration model can provide a good approximation to the SIR model when *R*_0_ < 1. The information criteria for this model was calculated with *p* = 2.

### Simulation experiments

In the following, we apply our theory and estimation methods to the SIR model. To generate data for estimation, we simulated time series of the number of individuals in each state according to our Markov process model using the Euler scheme of He *et al*. [20]. The pomp [29, 30] R package was used to implement the model. Our typical procedure was to simulate data with most of the parameters fixed at the default values in table 1 and for several choices of transmission rate.

The default parameters were chosen to be typical of an acute infectious disease of humans. Our default infectious period of 22 days is consistent with the combined latent and infectious period in a past model of pertussis [31], as is our default host mortality rate of 0.02 per year. Our default initial population size of 10 million is chosen to be similar to that of a very large city and to be large enough for the linear noise approximation to be reasonable. For setting the importation rate *η*, Keeling and Rohani [32] suggest using 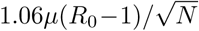 based on the time between extinctions and invasions of measles in England and Wales. Our own choice for the default *η* in table 1 is a simplification of this relation. The default observation frequency is chosen to be weekly because infectious disease notification data are often available at that frequency.

Simple sensitivity analyses of the distance estimates were carried out by allowing one or two of the parameters to vary from the default values. The full set of parameters used for each set of distance estimates is reported in the results. The initial values of the states were set to the equilibrium values and the model was run for 10 simulation years before sampling to allow the initially sampled states to vary according to the stationary distribution of the process. The sampling scheme was 1040 observations at a frequency of 1 observation per week. This corresponds to about 20 years of weekly observations, which is a realistic size for an epidemiological data set. Sampled time series of both the number infected and the number susceptible were used to generate an estimate of the distance to the threshold by the method described above. The true value for each estimate was calculated by plugging the simulation parameters into equation (8), solving for the fixed points, and calculating the eigenvalues of *F*. If there were two real eigenvalues, the informative eigenvalue was identified as the eigenvalue that would cross zero if the parameters were moved through the bifurcation point where *R*_0_ = 1 and *η* = 0.

We also conducted simulations with a linearly increasing transmission rate to evaluate the performance of estimates of changes in distance. To ease comparison with estimates from our other simulations, we used a similar amount of data for the individual distance estimates used to calculate the change in distance. The simulations were sampled for twice as long, the time series were split into two windows of 1040 weekly observations, and an estimate was obtained for each window.

### Software and reproducibility

Code to reproduce our results is available in an online repository [33].

## Results

### Main determinants of the accuracy of distance estimates

We first present some general considerations regarding when the distance to the epidemic threshold can be estimated from the fluctuation dynamics of the SIR model. Figure 1 shows representative examples of the kinds of time series that we suppose could become available for statistical analysis. For two of the parameter values, cycles are visible in both the number of susceptibles, *X*, and the number of infecteds, *Y*; which is a consequence of the eigenvalues of the Jacobian, *F*, being complex. This behaviour is typical of parameters for which *R*_0_ > 1. We have explained in Methods that, in this case, any white-noise perturbation may allow for the distance estimate to be obtained from either variable. When *R*_0_ < 1, typically there are two real eigenvalues. Without any knowledge about the model, we would expect that the ability to obtain an estimate depends on the covariance of the perturbations. With the knowledge that the observations come from an SIR model, we could expect that the dynamics of the *X* and *Y* variables will be largely independent of each other. The number infected will generally be too small to affect the fluctuations in the number susceptible. Thus the rate at which susceptible perturbations decay will depend mostly on the per-capita death rate *μ*, whereas the rate at which infected perturbations decay will depend on the sum of the per-capita rates at which *Y* grows and shrinks, *βX*\*/*N*_0_ – *γ* – *μ*. Thus the variable *Y* is generally the one that should be observed to estimate the distance to the threshold when the disease is not widespread. In the electronic supplementary material, we derive explicit equations for the autocorrelations that supports this conclusion.

**Figure 1:**
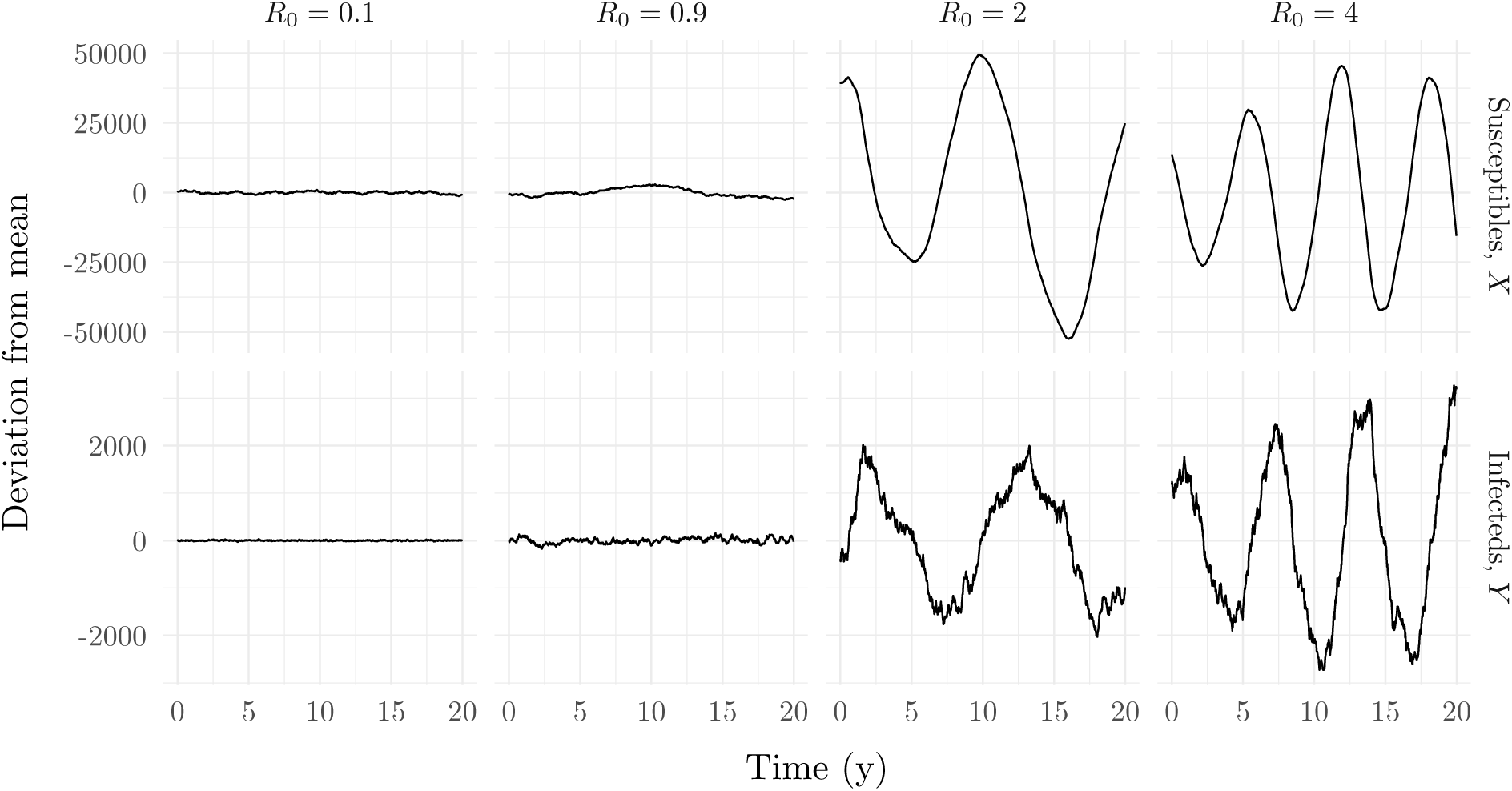
Example of simulated time series that could provide a distance estimate. Deviations are plotted instead of the simulated counts to align the time series vertically and because the deviations are what is used to calculate the estimate. The main idea is to estimate the rate at which deviations decay and oscillate. As these examples suggest, these rates tend to increase as the parameters of the system move away from *R*_0_ = 1. Parameters for the simulations are in table 1, with *β* set to *R*_0_(*γ* + *μ*).

Having provided some general insights into why distance estimates may be obtained from *Y* and not *X* when *R*_0_ < 1, we next consider a more specific answer for a specific set of parameters. We use the approach described in Methods to find the set of noise parameters that allow the distance to the threshold to be estimated with a given accuracy from each variable. These sets appear as regions in space in figure 2. Consistently with the conclusions of the previous paragraph, the regions are much larger for the number infected, *Y*, than for the number susceptible, *X*. The regions for *Y* include the perturbations that result from the intrinsic noise present in simulations of the model with finite population sizes. In contrast, a large part of the lower-error region identified for *X* is in fact not feasible because the covariance matrix constraint of 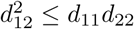 is not satisfied.

**Figure 2:**
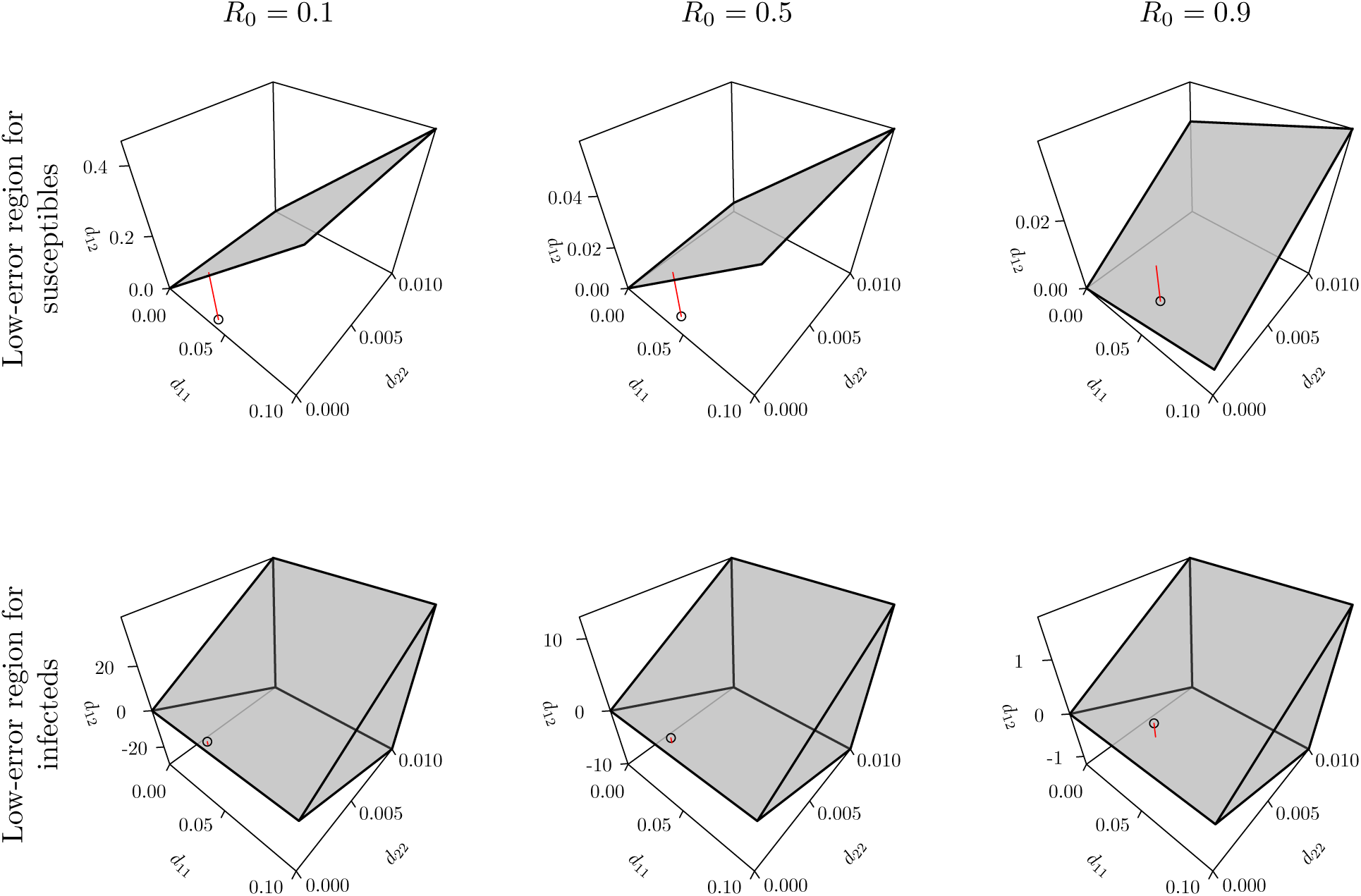
In our SIR model for most *R*_0_ < 1, the set of noise parameters under which the distance to the epidemic threshold may be estimated is much larger for the number infected than for the number susceptible. Each panel plots as a prism, for a range of variances in white-noise perturbations to the number susceptible and infected (subscripts 1 and 2, respectively) the region containing the covariances for the perturbations that result in the autocorrelation of the model variable being within 0.1 of the reference that would result in a perfect distance estimate. The axis labels denote elements of the matrix *D* in equation (7). The points in each panel correspond to the intrinsic perturbations calculated by the linear noise approximation equation (9). The red line segments are drawn vertically from these points to the nearest boundary of the low-error region. The points in the panels for the infecteds fall within the low-error region. Parameters were as in table 1, with *β* set to 1.66, 8.31 and 14.9.

Having shown that, in principle, distance estimates are often possible to obtain from the SIR model from at least one of the variables, we next turn to the question of whether estimates may be obtained in practice from a simulated time series of realistic length using our estimation method. Figure 3 shows that for time series of about 1000 observations our estimation method was generally successful when the perturbations are predicted to be suitable. The perturbations were simply intrinsic noise, so the low accuracy of estimates based on *X* when *R*_0_ = (0.1, 0.5, 0.9) is consistent with figure 2. As expected, when *R*_0_ > 1 estimates from both *X* and *Y* were similarly accurate. Therefore *Y* permits a distance estimate for all *R*_0_ considered. Fortunately, *Y* is the variable which is more often observed in practice. We have included estimates based on *X* in our results primarily to evaluate our analytical predictions about when a variable can provide an accurate distance estimate.

**Figure 3:**
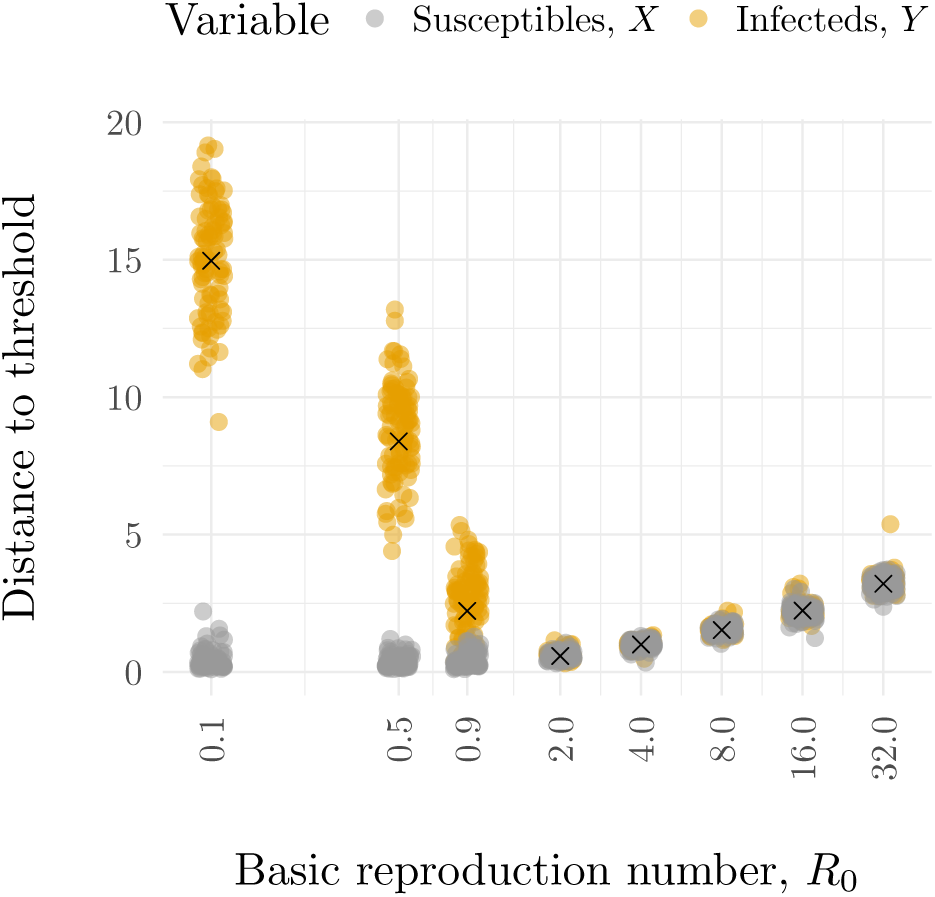
Distance estimates can be obtained from both *X* and *Y* in stochastic simulations of our SIR model. However, the distance estimates based on *X* are not always reliable. The *X* and *Y* variables were sampled weekly for 20 years. Parameters for the simulations are in table 1, with *β* set to *R*_0_(*γ* + *μ*). The distance to the threshold is defined here as the modulus of the informative eigenvalue. The true value is marked with an ‘x’. One hundred data sets were simulated for each set of parameters and all estimates obtained for each variable are plotted. The x-values of the points have been jittered to make the density of points at each y-value clearer.

### Effects of observation schemes on distance estimates

To begin evaluating how the observation scheme may affect estimates, we varied the frequency of observations in the time series. To avoid confounding the effects of frequency with the effects of the length of the time series, its length was kept the same for all sampling frequencies by adjusting the stop time of the simulations. In the *R*_0_ = 2 panel of figure 4, we see that when such a trade-off exists between the duration of observation and the observation frequency, a high observation frequency can be detrimental due to the short observation period. For daily observation frequency, the total duration of observation was limited to 1000 days, whereas the period length of the oscillations in the autocorrelation was about 11 years. With weekly observation frequency, the duration of observation is about 20 years and the estimates are much better. A rough guideline for accurate estimation seems to be that the duration of observation be at least as long as the period of any oscillation in the autocorrelation function. Another guideline is that the time between observations be much less than the period of oscillation of the autocorrelation. In the *R*_0_ = 16 panel of figure 4, no distance estimates where obtained when the observation frequency went from 0.25 per week to 1 per year. For these parameters, the autocorrelation had a period of about 3 years, so 3 observations per cycle seems much less likely to provide sufficient information than 40 observations per cycle. A similar guideline on the sampling frequency holds when the autocorrelation function is not periodic. In the *R*_0_ = 0.1 panel, no estimates based on *Y* are available as the sampling frequency dips below 1 per week. Here, the autocorrelation shrinks by a factor of e ≈ 2.7 about every 3.5 weeks. This time can be used to characterise the timescale of a decaying function and is sometimes called the return time. A third guideline, then, is that the time between samples should be less than the return time. In summary, for accurate estimates the duration of observation should be much greater than the time scale of the autocorrelation function but the time between observations should be much smaller than the time scale of the autocorrelation function.

**Figure 4:**
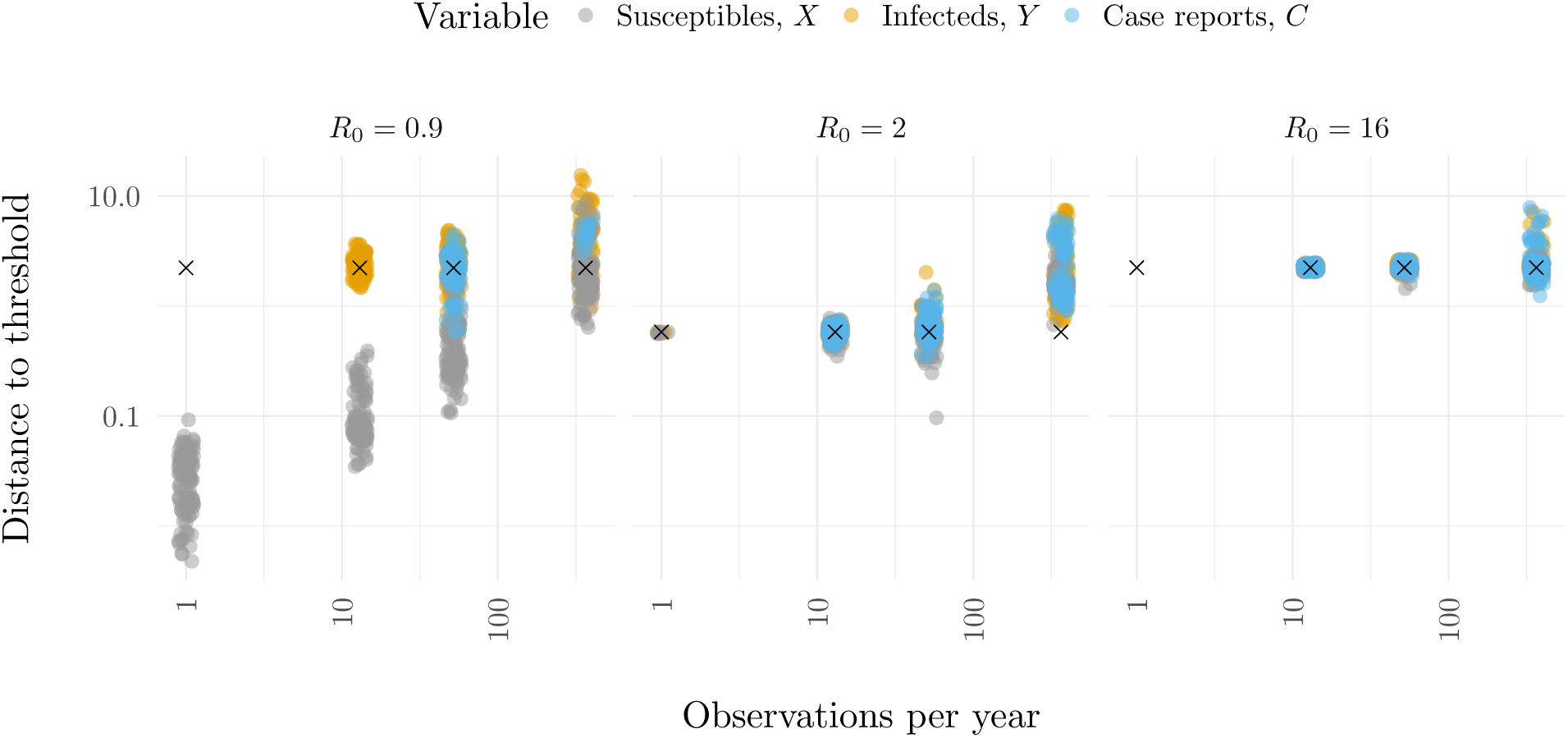
The availability of distance estimates is sensitive to the frequency of observations and whether the state of the time series consists of case reports. The true distance is marked with an ‘x’. The case reports variable is the accumulated number of recovery events since the last observation. Parameters besides the observation frequency are in table 1, with *β* set to *R*_0_(*γ* + *μ*).

In addition to sampling frequency, another key characteristic of observations is whether they represent direct observation of the state of the system or cumulative flows between states. In particular, it is relatively rare for the number of infections in a population at a given point in time to be observed. A more typical type of observation is the count of the number of infected individuals that moved into the removed class, for example because these individuals were diagnosed with infection and then greatly reduced contact with others [34]. We refer to such counts as case reports. Figure 4 shows that the estimates based on case reports were similar to those based on direct observation of *X* or *Y*. The main difference is that when the data consist of case reports we concluded more often that there was insufficient information available for an estimate. For example, many estimates based on *Y* are plotted when *R*_0_ = 0.9 and the observation frequency is 0.25 per week but the same set of simulations resulted in no estimates based on case reports. When the reporting of each recovery is not sure but instead occurs with a certain probability, as in the study of Gamado *et al*. [35], the number of simulations that resulted in estimates went down with the reporting probability (Fig. 5). In conclusion, estimates based on case reports can be as accurate as those based on state variables, but also can be less likely to be available for a given number of observations.

**Figure 5:**
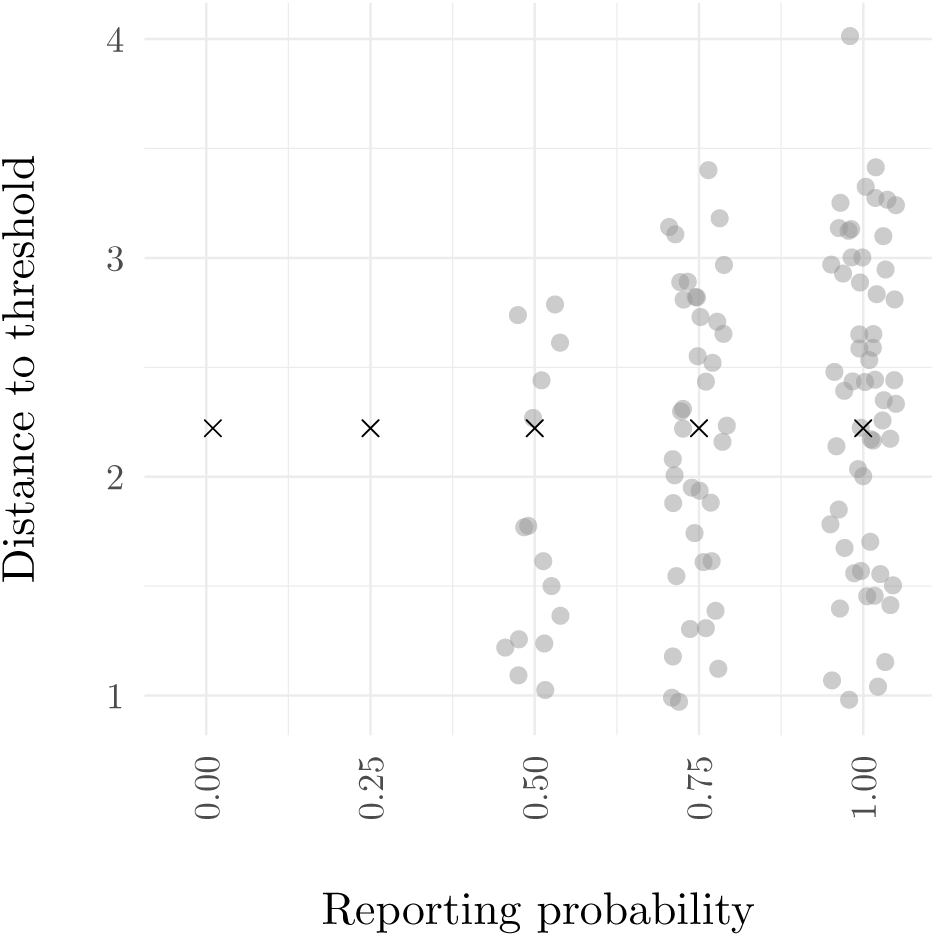
The accuracy of distance estimates did not clearly decline with the reporting probability but the number of times the data contained sufficient information for an estimate did. The true distance is marked with an ‘x’. These estimates were based on a time series of binomially distributed case reports, where the the accumulated number of recovery events since the last observation determined the number of trials and the reporting probability was the probability of success. Parameters besides the reporting probability are in table 1, with *β* set to 0.9(*γ* + *μ*).

The electronic supplementary material contains the results of our sensitivity analysis of environmental noise, population size, and our estimation of the rate of change of the distance to the threshold.

## Discussion

This work has presented a general solution to the problem of the selection of appropriate variables in multivariate systems for detection of slowing down as a threshold is approached. The solution is a method of calculating what type of white-noise perturbations, if any, allow slowing down to be detected based on observation of a given variable. To provide a specific example, this general solution has been applied to the SIR model and been shown to be consistent both with a model-specific analysis and with simulations. This application has also served to demonstrate and stress-test a method of estimating a distance to a threshold that is defined as one of the eigenvalues of the linearized model’s matrix *F*. Importantly, this informative eigenvalue is not always the dominant eigenvalue. When the informative eigenvalue is not dominant it is a consequence of the vital dynamics of the host occurring on a timescale that is much longer than the dynamics of small outbreaks that occur when the infection does not spread very well in the host population. Such a difference in time scales seems likely to occur in other multivariate models of population dynamics.

Looking beyond the SIR model, the question also arises of whether our method of identifying appropriate variables will be practical for models with many more than two degrees of freedom. In the SIR model, the autocorrelation function of one of the state variables is often very similar to that of one of the eigendirections. This allowed us to select variables based on the criteria of how well the autocorrelation function matched up with that of the eigendirection corresponding to the informative eigenvalue. In general, as the number of variables grows we might expect that the autocorrelation function of each variable to become more strongly influenced by multiple eigenvalues. For this more challenging case, we wonder whether harmonic inversion methods [36] might be capable of estimating the values of each of the eigenvalues that strongly influence each variable from its autocorrelation function. Variables that allow the informative eigenvalue to be estimated in this manner could then be considered appropriate for tracking the distance to the threshold.

The distance to thresholds in systems will generally change over time, and our results concluded with a simple demonstration of how these changes might be tracked. In the context of infectious disease surveillance, an exciting prospect of this approach is the possibility that surveillance programs might be able determine that some change in the system is moving it closer to the epidemic threshold long before the threshold is crossed. Besides increasing awareness, such measurement may allow for management of the distance to the threshold in some systems, for example by guiding the allocation of resources to vaccination programs. In this way, infectious disease control goals could move beyond early detection of and rapid response to epidemics toward targeted prevention of epidemics. Further, tracking has the potential to measure the relenting and reversing of system dynamics in response to control goals. Finally, establishing the conditions under which statistical analysis of fluctuations in the number of infected individuals is more informative than similar analysis of susceptible individuals does not make a case against susceptible reconstruction methods [37] in distance-to-threshold studies because such methods estimate major trends in the susceptible population size rather than fluctuations around them. Rather, our result makes a case for analysis of the fluctuations in the number infected, whether estimated from readily available time series incidence data or from pathogen sequence data [38].

## Authors’ contributions

EO conceived the main ideas in this paper. EO and JD designed the study. EO performed simulations and analysis. All authors participated in the interpretation of results, drafting of the manuscript and gave final approval for publication.

